# Machine learning for lumbar and pelvis kinematics clustering

**DOI:** 10.1101/2022.05.23.493131

**Authors:** Seth Higgins, Sandipan Dutta, Rumit Singh Kakar

**Affiliations:** Human Movement Science, Oakland University, Rochester Hills, MI, USA; Mathematics and Statistics, Old Dominion University, Norfolk, VA, USA

**Keywords:** machine learning, clustering, lumbar, pelvis, kinematics

## Abstract

Clustering algorithms such as k-means and agglomerative hierarchical clustering (HCA) may provide a unique opportunity to analyze time-series kinematic data. Here we present an approach for determining number of clusters and which clustering algorithm to use on time-series lumbar and pelvis kinematic data. Cluster evaluation measures such as silhouette coefficient, elbow method, Dunn Index, and gap statistic were used to evaluate the quality of decision making. The result show that multiple clustering evaluation methods should be used to determine the ideal number of clusters and algorithm suitable for clustering time-series data for each dataset being analyzed.

## Introduction

Traditional statistical methods comparing movement kinematics in biomechanics involve calculating discrete variables like peak joint positions or ranges of motion and comparing between pre-defined groups based on age, sex, pathology, or condition. However, this method generalizes that the movement patterns of individuals within a group are similar and inherently different from those of the group being compared to, which might not be the case. A similar approach is commonly used when studying lumbopelvic or trunk kinematics where peak joint positions and ranges of motion of the spine are compared to predetermined groups (i.e., low back pain vs. no low back pain; old vs. young) during trunk flexion-extension movements (Paquet et al. 1994; Wong and Lee 2004; Pries et al. 2015; Laird et al. 2019). Past research has found that individuals with low back pain have reduced lumbar spine and pelvis range of motion during trunk flexion (Paquet et al. 1994; Wong and Lee 2004; Laird et al. 2019) and older adults (50-75 years) have reduced lumbar spine range of flexion and greater pelvis range of flexion during trunk flexion when compared to younger adults (Pries et al. 2015).

Analyzing the entire time-series data instead or in-addition to discrete variables may be useful for lumbopelvic kinematics and to answer other biomechanical questions. Time-series analysis using machine learning techniques such as clustering offers a convenient method to distinguish movement patterns within a group of individuals. Clustering aims to take an unlabeled set of data and organize them into homogenous groups (Warren Liao 2005). Clustering may help separate lumbar spine and pelvis kinematics into multiple distinct movement patterns, which may help researchers identify different lumbar spine and pelvis movement patterns within a heterogenous group of healthy young and old adults and individuals with low back pain. We use lumbar and pelvic kinematics here to show the applicability of clustering analysis in biomechanics in healthy adults.

Clustering approaches have been used previously in biomechanics research to segment kinematic and kinetic data (Sawacha et al. 2010; Bennetts et al. 2013; Ackermans et al. 2019; Sawacha et al. 2020). However, these studies focus on specific events and discrete variables extracted from a timeseries instead of the entire time-series (Sawacha et al. 2010; Bennetts et al. 2013; Ackermans et al. 2019). While other studies with a focus on time-series have used clustering algorithms using standard or squared Euclidean as the distance metric (Phinyomark et al. 2015; Sawacha et al. 2020) to measure the similarity or dissimilarity among data points (Singh et al. 2013). Euclidean distance is a correlation-based distance metric that calculates the root of square difference between the coordinates of a pair of data points (Singh et al. 2013; Aghabozorgi et al. 2015). Along with requiring the two time-series being compared to be the same length, which is often not common in joint kinematics, Euclidean distance is also sensitive to small shifts or delays in time (Kate 2016). Thus, even though Euclidean distance is simple and efficient, it may not be the most appropriate for calculating the similarity in joint kinematics data. Euclidean distance is generally performed on time-series data that has been transformed using Fourier transforms, wavelets, or Piecewise Aggregate Approximation (Aghabozorgi et al. 2015). Within raw time-series data where temporal drift is common, shape-based distance metrics may be useful because the time of occurrence is not important when calculating the distance between time-series. In these situations, elastic methods such as dynamic time warping (DTW) are used rather than Euclidean distance (Kate 2016; Lee 2019). For example, DTW has been previously used in speech recognition to match words even when timing and pronunciation varies from person to person, and also to analyze the similarity of gait patterns (Berndt and Clifford 1994; Adistambha et al. 2008; Lee 2019).

As the amount of data in biomechanics and the interest in machine learning increases, we need to determine best practices to ensure accurate analysis and reproducibility (Halilaj et al. 2018). There are many different clustering techniques to choose from. K-means and agglomerative hierarchical clustering (HCA) are commonly preferred for time-series data analysis (Sawacha et al. 2010; Bennetts et al. 2013; Ackermans et al. 2019). Each algorithm has its strengths and weaknesses, which makes it difficult to choose the most optimal one for a given situation. For example, k-means is computationally faster than HCA and can handle many variables, but the number of clusters need to be predefined, and the results are dependent on the initial centroid definition (Dhanachandra 2015; Aghabozorgi et al. 2015). On the other hand, HCA does not require any predefined number of clusters, but this method is more computationally expensive (Bouguettaya et al. 2015). To select the most optimal algorithm for a dataset, different clustering approaches might need to be compared. Therefore, the purpose of this study was to compare clustering performance of the k-means and HCA algorithms for classifying lumbar and pelvis kinematics during trunk flexion and extension task.

## Methods

### Participants

Eighty-four healthy adults completed this study (Table 1). Participants were split up into two age groups: young (18-40 years) and middle–age (41-65 years) adults, including both male and female participants. Healthy adults were free from any medical condition/injury that would interfere with participation, had not completed a major competitive event one week before the data collection, and completed a pre-participation health status, a medical history questionnaire, and a short-form International Physical Activity Questionnaire to assess eligibility.

**Table 1:** Participant Characteristics.

|  | Young Adults (n = 50) | Middle-age Adults (n = 34) |
| --- | --- | --- |
| Age (years) |  |  |
| (age range; mean±SD) | 18-40; 26.84±6.77 | 41-65; 52.23±6.89 |
| Mass (Kg) | 67.17±13.4 | 70.27±11.22 |
| Height (m) | 1.69±0.08 m | 1.68±0.07 m |
| Physical activity duration |  |  |
| (hours/ week) | 6.2±3.3 | 8.1±8.7 |
| Males | n = 14 | n = 17 |
| Females | n = 36 | n = 17 |

### Data Collection and Analysis

Informed consent was obtained prior to data collection, as approved by the university institutional review board. This was a secondary analysis from a larger research project with the focus on the kinematics of the trunk and pelvis captured in the sagittal plane during 3 trials using a reflective marker-based motion capture system (Sampling frequency: 120 Hz; Vicon^®^ Motion System Ltd, Oxford, UK). The marker setup has been described in detail previously (Li et al. 2017; Kakar et al. 2018). In brief, the reflective markers were placed on the jugular notch, xiphoid process, spinal process of the C7, T2, T4, T6 and T8, spinal process of the T10 and T12, right and left transverse processes of T11, spinal process of the L2 and L4, right and left transverse processes of L3, and ASIS and PSIS on the right and left (Figure 1). The lumbar was constructed using the spinal process of the L2 and L4, and right and left transverse process of L3. The pelvis was constructed with CODA model (Bell et al 1989). The segmental angles of the lumbar spine and pelvis were calculated relative to the global coordinate system. The following x-y-z Cardan rotation sequence was used: extension (+)/flexion [or pelvic posterior (+)/anterior tilt].

**Figure 1:**
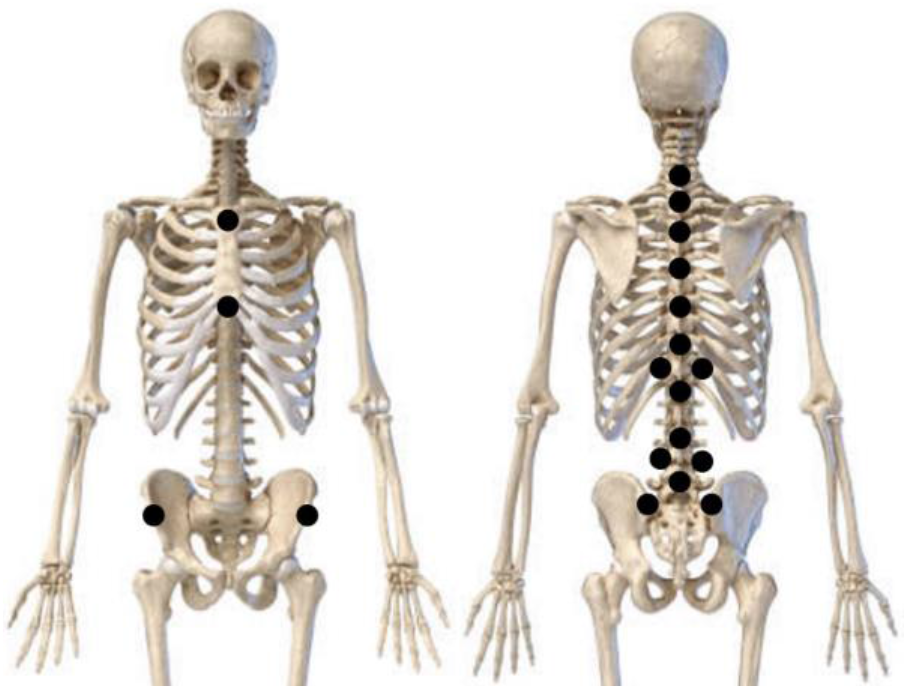
Marker placement on the trunk and pelvis.

Participants began each trial in neutral standing position, moved into their maximal trunk flexion range (max flexion), and then returned to neutral/ starting position (neutral). Participants continued the motion into maximal trunk extension (hyperextension) before returning back to neutral to end the trial. The entire movement task was performed as one fluid motion. Lumbar and pelvis kinematics were calculated using Visual 3DTM software (V.5; C-Motion Inc., Maryland, USA).

### Clustering Analysis

#### K-means Clustering

Using k-means clustering via a custom MATLAB code (GitHub), we investigated the implementation of grouping lumbar spine and pelvis kinematics of young and middle-aged adults during a trunk flexion/extension task. K-means is a partitioning clustering algorithm (k = number of clusters/ groups), first proposed in 1967 (MacQueen 1967). K-means aims to minimize the total distance between all objects in a cluster from their cluster center (Warren Liao 2005; Aghabozorgi et al. 2015). A cluster is represented by its centroid, which is a mean of the time-series within a cluster (Rai and Singh 2010). K-means starts by first determining the number of clusters and centroids (Warren Liao 2005; Aghabozorgi et al. 2015; Phinyomark et al. 2018; Halilaj et al. 2018). Each object is then assigned to the cluster with the closest centroid, which minimizes the overall within-cluster dispersion (Abbas 2008; Rai and Singh 2010). DTW was the distance metric used to assign each time-series to the closest centroid. The previous centroids are then replaced by the newly formed clusters and the algorithm repeats this process until no objects change clusters (Warren Liao 2005). Execution of k-means is described in figure 2.

**Figure 2:**
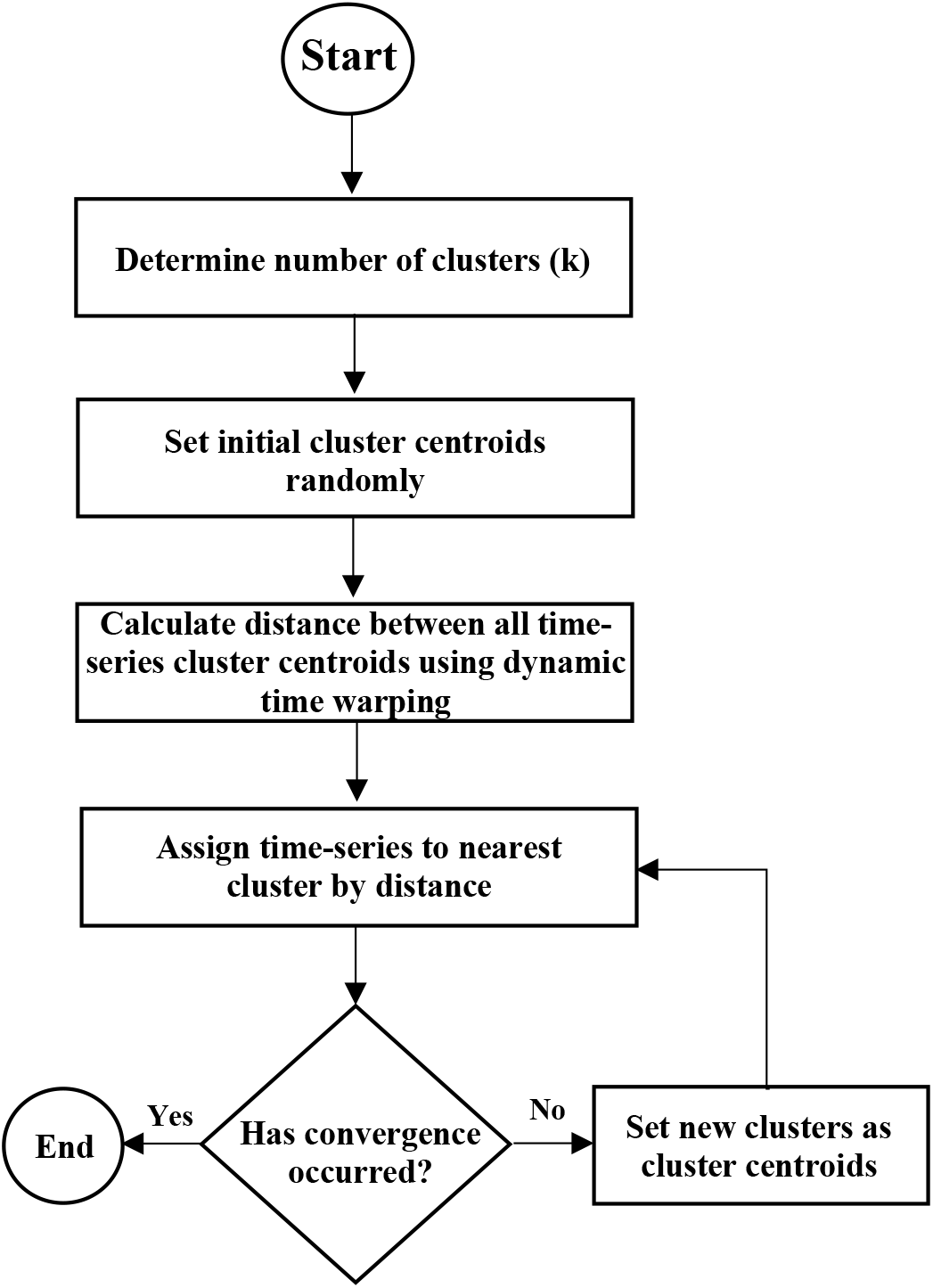
Flow diagram of k-means clustering procedure.

#### Hierarchical Clustering (HCA)

HCA is a “bottom-up” approach that considers each time-series as an individual cluster, and then gradually combines the clusters until all time-series are grouped in one cluster (Aghabozorgi et al. 2015). The equation below is for HCA:

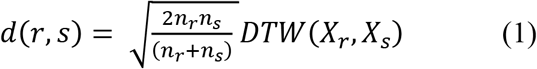

Where, DTW (X_r_, X_s_) is the DTW distance between X_r_ and X_s_, which are the centroids of clusters r and s and n_r_ and n_s_ are the number of elements in clusters r and s.(Equation 1). HCA starts off by calculating the distance between every time-series of all participants using DTW. Second, individual clusters are combined into binary clusters using the Ward linkage method (Ward Jr 1963). The aim of Ward linkage is to minimize the total within-cluster variance by merging clusters that lead to a minimum increase in total within-cluster variance post-merger (Warren Liao 2005; Charrad et al. 2014). Lastly, newly formed clusters are grouped into larger clusters until the dendrogram is formed (Phinyomark et al. 2015). The method for calculating HCA is represented in figure 3.

**Figure 3:**
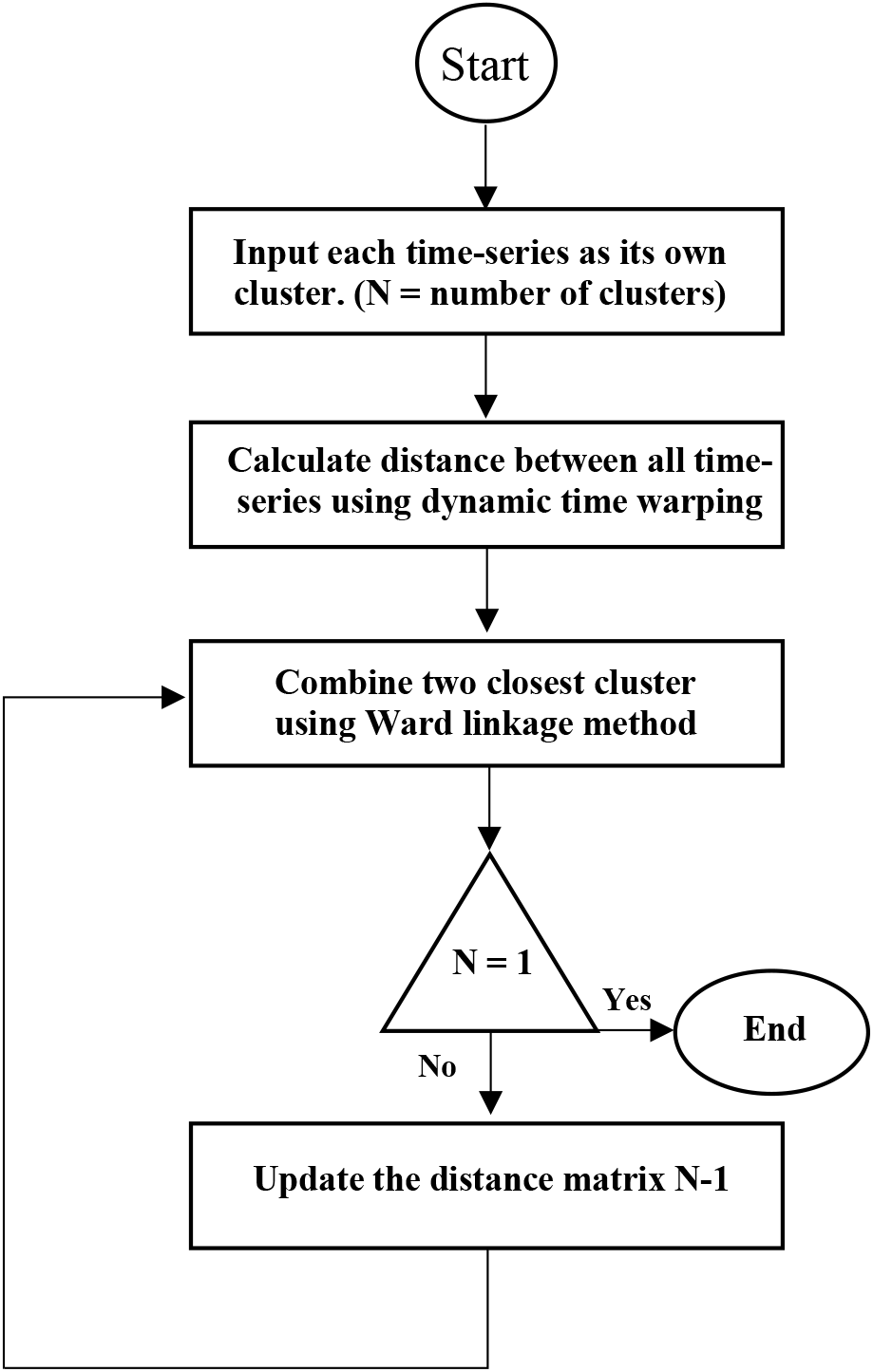
Flow diagram of HCA procedure.

### Distance Measures

#### Dynamic Time Warping (DTW)

DTW is a shape-based distance metric that measures the similarity between two timeseries, which may vary in speed (Aghabozorgi et al. 2015). DTW starts out with two time-series (Q and C) (Equation 2). Then the distance is computed by first finding the best alignment between them by creating a n-by-m matrix whose (*i^th^, j^th^*) element is equal to (*q*_i_ – *c*_j_)^2^ which represents the cost to align the point *q*_i_ of the time-series Q with the point *c*_j_ of time-series C (Kate 2016). An alignment between the two time-series is represented by a warping path, W = *w*_1_, *w*_2_, …, *w_k_*, …, *w_K_*, in the matrix (Kate 2016). The best alignment is the path that minimizes the total cost of aligning its points, which is termed as the DTW distance (Kate 2016).

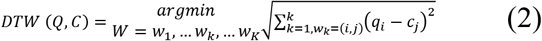

The warping path was found using dynamic programming algorithm (Berndt and Clifford 1994; Keogh et al. 2004).

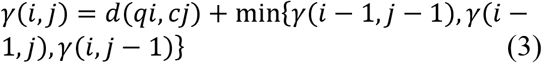

Where *d*(*i,j*) is the distance found in the current cell, and *γ*(*i,j*) is the cumulative distance of *d*(*i,j*) and the minimum cumulative distance from the three adjacent cells (Keogh et al. 2004) (Equation 3).

#### Clustering Evaluation

Four independent evaluation measures were used to compare k-means and HCA performance using two methods. For the first method, clustering evaluation measures were used to determine the optimal number of clusters used for both kinematic datasets and for both clustering methods (Rousseeuw 1987; Tibshirani et al. 2001; Kryszczuk and Hurley 2010; Zhou et al. 2017; Syakur et al. 2018; Yuan and Yang 2019). The optimal number of clusters were determined by calculating each evaluation measure for different number of clusters (k = 2-10) (Saputra et al. 2020). For each fixed value of k, k-means was ran 30 times and each evaluation measure was calculated (Wang et al. 2017). The criteria for determining the number of clusters for each evaluation measures is described below. For the second method, each evaluation measure was used to compare the quality of the clusters between k-means and HCA (Xu et al. 2006).

#### Silhouette Coefficient

Silhouette coefficient was first developed in 1987 as a graphical aid to help interpret and validate clustering analyses (Rousseeuw 1987). Silhouette coefficient measures how similar a time-series is to the time-series within its own cluster compared to the timeseries in other clusters (Sawacha et al. 2010). To calculate silhouette coefficient, the average distance of each time-series to all the other time-series within the same cluster was calculated (a). Then, the average distance of each time-series to all the other time-series in the nearest cluster by distance was calculated (b). DTW was used as the distance metric (Berndt and Clifford 1994; Kate 2016). The values of a and b for each time-series were used to calculate the silhouette coefficient using the equation below:

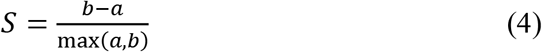

Silhouette coefficient ranges from −1 to +1, and −1 indicates that the time-series was assigned to the wrong time-series, 0 indicates that the time-series could be assigned to either cluster, and +1 indicates that the time-series was assigned to the correct cluster. The silhouette coefficients for each time-series were averaged to get an overall silhouette score. The number of clusters (k) that produced the greatest silhouette score indicated a higher quality result and was chosen as the k for further analysis (Xu et al. 2006; Jun and Lee 2010; Zhou et al. 2017; Navarro and Moreno-Ger 2018; Yuan and Yang 2019; Sebastian et al. 2020). When comparing the silhouette scores between the two clustering methods, the greatest silhouette score indicated the clustering method that produced the highest quality clusters.

#### Elbow Method

The elbow method was another clustering evaluation technique used to determine the optimal number of clusters on a set of data. For each number of clusters (k), the total sum of square error was calculated and then plotted. One drawback of this method is that when the k value increases, the total sum of square error value will reach near 0, which will not determine if a cluster is good or bad based on the measurement alone (Saputra et al. 2020). With this drawback in mind, this method suggests that the best value of k is when the total sum of square error value reduces significantly, or the “elbow” of the graph (Syakur et al. 2018). When comparing the elbow method results between k-means and HCA, the lowest total sum of square error indicated the clustering method that produced the highest quality clusters.

#### Gap Statistic

Gap statistic is a clustering evaluation technique developed in 2001 to provide a statistical procedure to determine the optimal number of clusters (Tibshirani et al. 2001). To calculate the gap statistic, the data is clustered into k clusters. Let *d_ii_*’ be the distance between time-series *i* and each centroid *i*’. DTW was used as the distance measure. In equation 5, Dr is the sum of the distances for all the time-series in cluster r (C_r_).

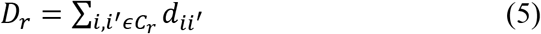

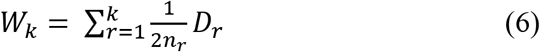

In equation 6, W_k_ is the pooled within-cluster sum of squares for each cluster, where D_r_ is the sum of the distances for all time-series in cluster r and n_r_ are the number of elements in cluster r. Next, the log(W_k_) was compared with its expectation under a null reference distribution of the data. This is defined as 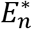, which is the expectation under a sample of size n from the reference data. The estimated number of clusters (k) was the value maximizing Gap_n_(k) (Equation 7).

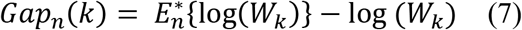

#### Dunn Index

Dunn index is a clustering evaluation algorithm developed in 1974 (Dunn 1974). Dunn index is defined by equation 8 where the min is the minimum distance between any two time-series that fit in different clusters, and max is the maximum distance between two time-series in the same cluster (Dunn 1973). δ(C_i_,C_j_) is the inter-cluster distance between cluster C_i_ and C_j_. Δ(C_k_) is the intracluster distance within cluster Ck (Equation 8). The higher Dunn index value indicates the optimal number of clusters for a given data set and the optimal clustering solution when comparing different clustering algorithms (Kryszczuk and Hurley 2010).

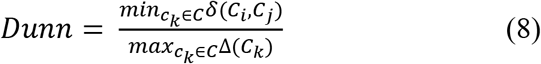

## Results

Cluster evaluation results for k-means and HCA are represented in figure 4. Based on the criteria for each evaluation measure, the optimal number of clusters for each clustering algorithm and dataset were determined (Table 2). Since the optimal number of clusters changed with the choice of the evaluation measure, a list of the top 3 choices for the number of clusters were made for each of the four evaluation measures (Table 2). Based on majority ranking (Sikdar et al. 2020), there was a clear preference of 4 clusters for the lumbar kinematics using HCA and 2 clusters for the pelvis kinematics using HCA and k-means. The evaluation results for lumbar kinematics using k-means were less consistent. Therefore, to compare HCA results with k-means, we selected 4 for k-means as well, since it was also amongst the top choices. The number of participants within each cluster are represented in table 3.

**Figure 4:**
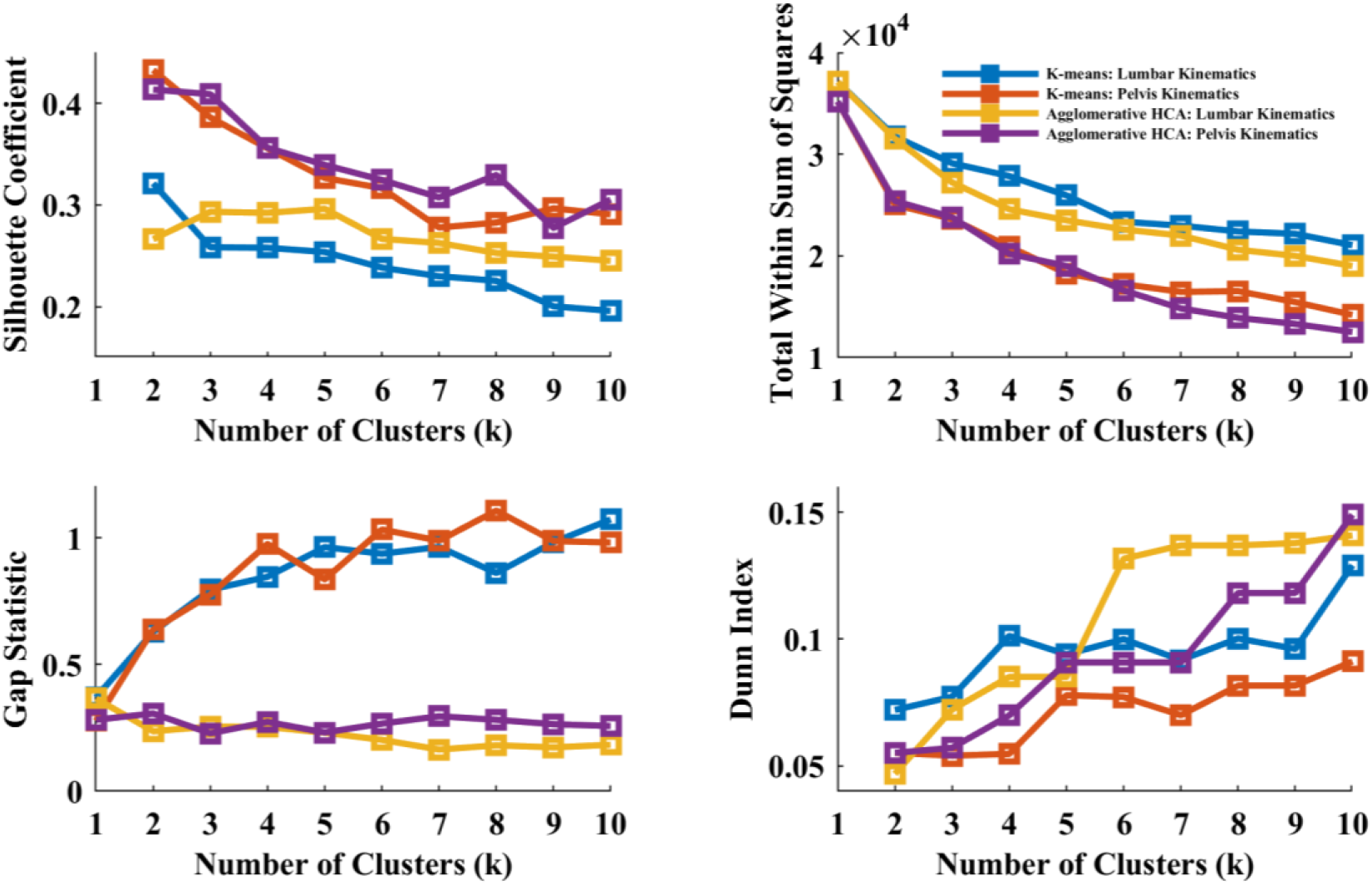
Cluster evaluation results for k-means and HCA.

**Figure 5:**
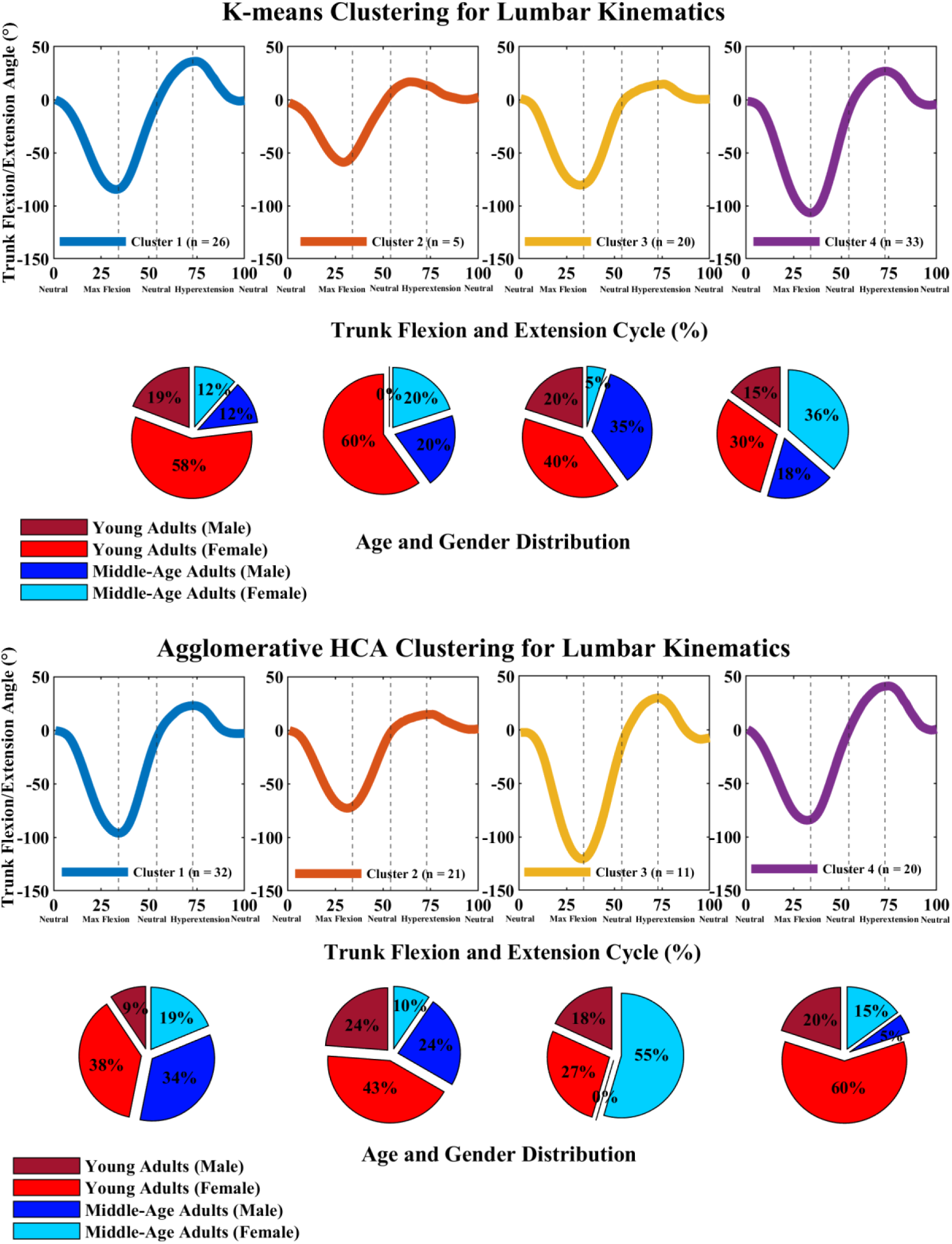
K-means and HCA clustering results for lumbar kinematics during trunk flexion/extension task.

**Figure 6:**
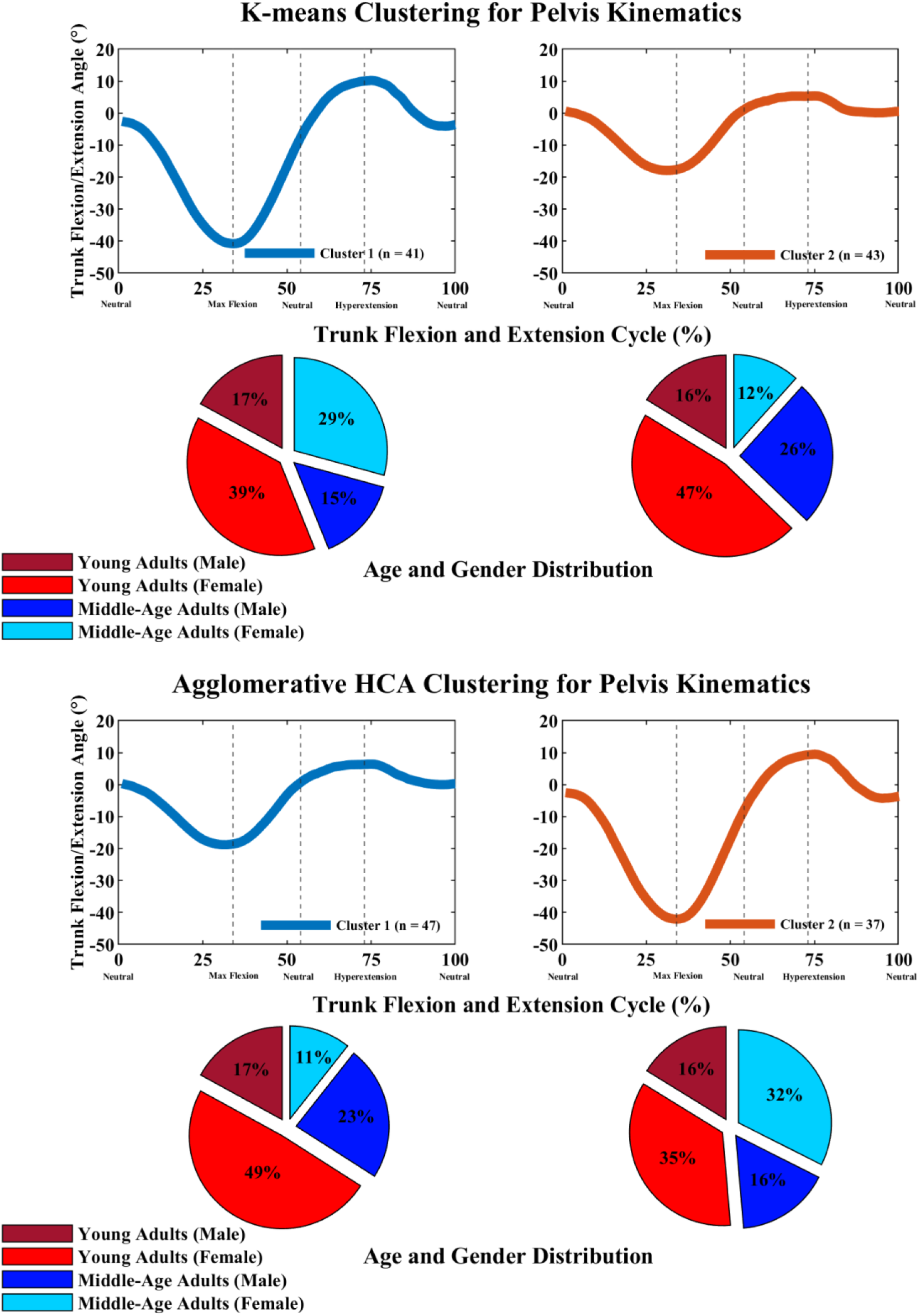
K-means and HCA clustering results for pelvis kinematics during trunk flexion/extension task.

**Figure 7:**
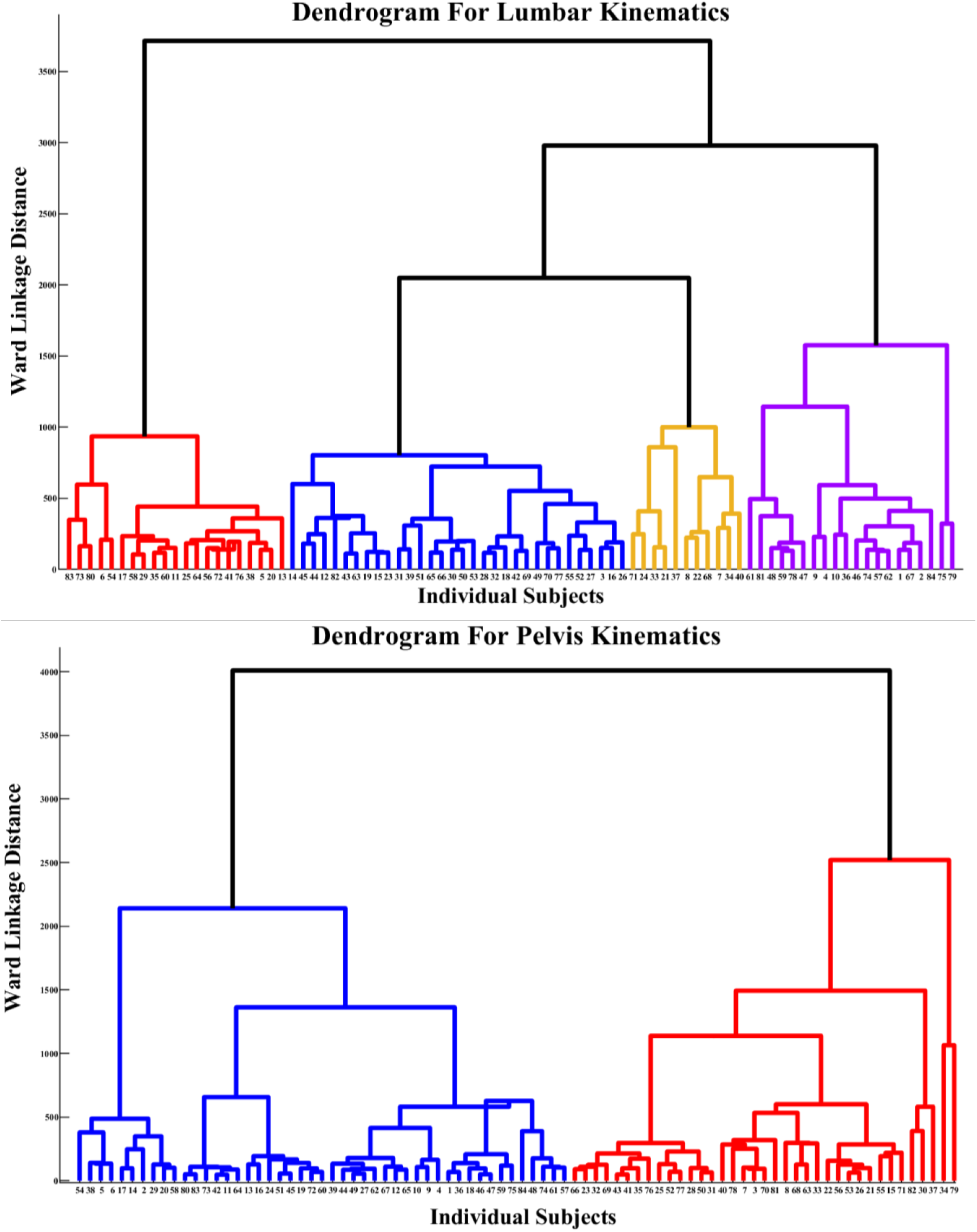
Ward minimum variance linkage dendrogram for the HCA cluster analysis of the lumbar (top) and pelvis (bottom) kinematics during trunk flexion/extension task. For lumbar dendrogram, four groups are highlighted for cluster 1 (blue), cluster 2 (red), cluster 3 (yellow), and cluster 4 (purple). For pelvis dendrogram, two groups are highlighted for cluster 1 (blue) and cluster 2 (red). Participant numbers are indicated in the x axis. Participant numbers are indicated in the x axis.

**Table 2.** Top three choices for optimal number of clusters and their validation measure scores (k; score) for each clustering algorithm and each validation measure. Values are in order of their ranking with the first value being the choice of k that produces the optimal value of the validation measure. Bolded* cells represent the optimal number of clusters (k) selected for that clustering algorithm and dataset.

|  | Validation Measures | Silhouette Coefficient | Elbow Method | Gap Statistic | Dunn Index |
| --- | --- | --- | --- | --- | --- |
| K-means | Lumbar | 2; 0.321 | 6; 23,355 | 10; 1.07 | 10; 0.129 |
|  |  | 3; 0.258 | 7; 22,975 | 9; 0.982 | **4\*; 0.101** |
|  |  | **4\*; 0.258** | 8; 22,401 | 7; 0.964 | 8; 0.100 |
|  | Pelvis | **2\*; 0.433** | **2\*; 25,038** | 8; 1.107 | 10; 0.091 |
|  |  | 3; 0.386 | 3; 23,627 | 6; 1.033 | 8; 0.082 |
|  |  | 4; 0.356 | 4; 20,899 | 7; 0.988 | 9; 0.082 |
| HCA | Lumbar | 5; 0.296 | **4\*; 24,587** | 1; 0.366 | 10; 0.141 |
|  |  | 3; 0.293 | 5; 23,485 | 3; 0.255 | 9; 0.138 |
|  |  | **4\*; 0.292** | 6; 22,556 | **4\*; 0.254** | 8; 0.137 |
|  | Pelvis | **2\*; 0.414** | **2\*; 25,359** | **2\*; 0.307** | 10; 0.149 |
|  |  | 3; 0.409 | 3; 23,808 | 7; 0.296 | 9; 0.118 |
|  |  | 4; 0.356 | 4; 20,216 | 8; 0.282 | 8; 0.118 |

**Table 3:**
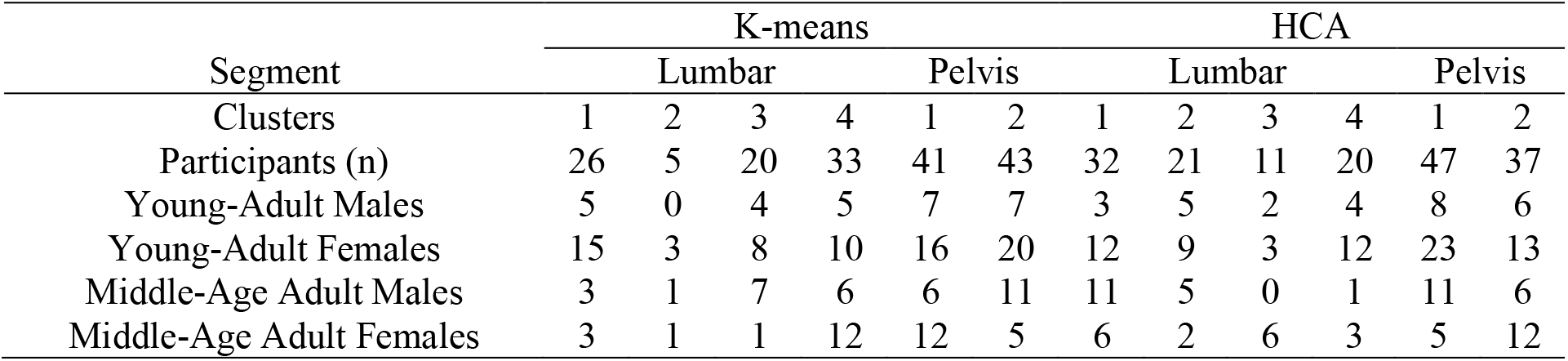
Breakdown of the number of participants and number of young adult and middle age adult males and females within each cluster for each clustering algorithm and dataset.

Using the number of clusters determined above, all four clustering evaluation measures were compared between k-means and HCA for both datasets to determine which clustering algorithm produced the best clustering results. For the lumbar kinematics HCA produced a greater silhouette coefficient (HCA: 0.292; k-means: 0.258) and lower total within sum of squares for the elbow method (HCA: 24,587; k-means: 27,852) than k-means, but k-means produced better clustering results based on the results of the Dunn Index (HCA: 0.085; k-means: 0.101) and gap statistic (HCA: 0.254; k-means: 0.846). HCA was chosen as the most optimal clustering algorithm for the lumbar kinematic data because HCA was more consistent when considering the majority ranking results. For the pelvis kinematics, k-means was chosen as the most optimal clustering method because k-means produced better clustering results when examining the evaluation results from silhouette coefficient (HCA: 0.414; k-means: 0.433), elbow method (HCA: 25,359; k-means: 25,038), and gap statistic (HCA: 0.307; k-means: 0.638), but for the Dunn Index (HCA: 0.055; k-means: 0.055), k-means and HCA produced the same value.

## Discussion

This study aimed to compare clustering performance of k-means and HCA clustering algorithms for time-series kinematic data using lumbar and pelvis segmental kinematics during lumbar flexion and extension motion. The current study was performed to showcase that machine learning and statistical approaches such as clustering algorithms may be better suited to categorize and compare population groups based on time-series analysis of the joint kinematics instead of solely relying on comparing discrete kinematic variables between predetermined groups.

The clusters were able to identify time series with differences in the degree and timing of peak lumbar and pelvis flexion and extension and was not restricted by categorization of participants based on age group and sex demographics. Based on the amount of motion, cluster #2 from HCA, was characterized by the smallest degree of lumbar flexion and extension, and included more young adults (n = 14) than middle-age adults (n = 7) and a similar amount of females (n = 11) and males (n = 10), while cluster #3, included individuals with the greatest degree of lumbar flexion, constituted a similar amount of young (n = 5) and middle-age adults (n = 6), and more females (n = 9) than males (n = 2). For the pelvis kinematics clusters from k-means, there are more young adults (n = 23) and less middle-age adults (n = 18), and more females (n = 28) than males (n = 13) in the cluster with the greatest anterior pelvic tilt (cluster 1) and more young adults (n = 27) and less middle-age adults (n = 16) and more females (n = 25) than males (n = 18) in the cluster with the least anterior pelvic tilt (cluster 2).

Past research has reported that older adults (51-75 years old) have reduced lumbar spine flexion range and greater pelvis anterior tilt during trunk flexion when compared to younger adults (Pries et al. 2015). Results of the current study show that reduced lumbar flexion may not be entirely age related. Since both young and middle-age adults were represented within each cluster, it cannot be assumed that an individual will have greater/ reduced lumbar flexion and extension based on age, but rather an individual characteristic.

Populations with low back pain and spinal fusions for idiopathic scoliosis have also been characterized as having reduced lumbar range of motion and lumbopelvic rhythm (Paquet et al. 1994; Wong and Lee 2004; Wilk et al. 2006; Laird et al. 2019). Utilizing clustering results of normative lumbar and pelvis kinematics may help us better understand movement adaptations and variations in individuals with low back pain and spinal fusions for idiopathic scoliosis. By comparing an individual’s performance to normative movement groups (clusters here), identifying where the phase lags and hypermobility’s are may help physicians and physical therapist develop personalized rehabilitation strategies to treat these individuals.

From a methodological perspective, the results of this study show that when selection of the most optimal algorithm or the number of clusters for each clustering algorithm, multiple evaluation measures can be used. Since each evaluation measure produced different results within the same algorithm and dataset, it is not sufficient to rely on a single evaluation measure and compare results across algorithms. Paired with evaluation measures, a rank-based approach, was critical in determining the number of clusters (Sikdar et al. 2020) in the current study. This approach determined that the optimal number of clusters as 4 for lumbar kinematics and 2 clusters for pelvis kinematics, which also matches with the common approach of physiological classification of the participants into young- and middle -aged males and females.

Overall, there are many things to consider when using clustering algorithms. First, each clustering algorithm takes a different approach. K-means clusters data by grouping all the data within the centroid that is closest by distance (DTW or Euclidean). HCA clusters data by merging the two closest data points together in a hierarchical fashion until all the data is merged into one cluster. Second, there are different distance measures (example, DTW or Euclidean) that can be used. Euclidean distance is generally used as the distance metric for most clustering algorithms, but it may not be appropriate for calculating the distance between two timeseries. Along with requiring the two timeseries to be the same length, Euclidean distance is also sensitive to even small shifts or delays in time (Kate 2016). DTW was used in this study because this distance metric was developed to overcome the limitations of Euclidean distance metric (Kate 2016; Lee 2019). Third, there are different clustering evaluation methods to choose from. This study used silhouette coefficient, elbow method, Gap Statistic, and Dunn Index to evaluate clustering results, but other clustering evaluation metrics such as Calinski Harabaz Index, Davies Bouldin Index can be used (Calinski and Harabasz 1974; Davies and Bouldin 1979).

There are several limitations present in this study. First, there are many clustering algorithms and validation measures available, but only 2 commonly used algorithms were compared through 4 different outcome measures. Although, the evaluation process presented in this study can be generalized to other clustering methods and validation measures. Second, we only tested young and middle-age adults. Further research on this topic should be conducted on older adults (age > 65), and populations with musculoskeletal pathologies directly effecting low-back, such as mechanical low back pain. Third, we only showcased one use-case of the clustering approach for timeseries analysis through trunk flexion/extension task, while these methods can be applied to other time-series data, both kinematics and kinetics, and during various movement tasks. Lastly, the number of individuals in the two different age and sex groups were not the same. The current study was not aimed at comparing groups directly, but rather categorizing individuals based on their movement performance. To simultaneously compare group differences, a more even distribution of the participants in each group may be useful.

In conclusion, this study proposed categorizing time-series data such as segmental/ joint kinematics (lumbar spine and pelvis kinematics in this study), to compare movement differences through different clustering algorithms, k-means, and HCA. Additionally, evaluation measures are important for the decision-making process of selection of optimal number of clusters, and multiple evaluation measures should be used simultaneously rather than relying on one, since results can vary based on the length, and other characteristics of a time-series. Rank-based approach was found to be useful when comparing results of different evaluation measures, and for eventual selection of optimal clusters based on scores and frequency.

## Acknowledgments

The authors would like to thank all the individuals who volunteered their time for participation in this research project. Authors would also like to thank the undergraduate students that helped at various stages of the duration of this project.

